# Benchmarking three simple DNA staining-based image metrics for live-cell tracking of chromatin organization

**DOI:** 10.64898/2026.03.30.715467

**Authors:** Minwoo Kang, Aidan Tomas Cabral, Manasi Sawant, Hawa Racine Thiam

## Abstract

Quantifying chromatin-state dynamics in living cells remains challenging, in part because most methods require fixation or cell lysis. Here, we benchmark and introduce three simple live-cell image-derived metrics computed from routine DNA staining—the coefficient of variation (CV), 1-Gini, and the Diffuse Signal Index (DSI), introduced here—as fixation-free readouts of chromatin organization. Using HL60-derived neutrophils (dHL-60) undergoing NETosis as a model system with a pronounced compact-to-decompact chromatin transition, we show that all three metrics track progressive chromatin reorganization in live-cell trajectories, with DSI providing the strongest trajectory-level discrimination between NETing and non-NETing cells. Comparison with Tn5-based chromatin accessibility measurements in fixed cells shows that all three metrics correlate with chromatin accessibility, supporting their biological interpretability. We further show that all three metrics track chromatin reorganization in a second, mechanistically distinct biological process, capturing mitotic chromatin compaction and post-mitotic chromatin decompaction in U2OS cells. Together, our results provide a practical framework for extracting chromatin-organization readouts from routine live-cell DNA staining and highlight that metric performance is task- and context-dependent. To make our metrics accessible to a broad audience, we built NucMetrics, an open-source ImageJ/Fiji macro toolset that computes all three metrics without requiring programming.

## Introduction

Chromatin organization is dynamically remodeled during cell-state transitions^1–7^, and monitoring these changes in living cells is important to understand how chromatin dynamics regulates nuclear architecture and cell physiology^8,9^, as well as how dysregulated chromatin organization alters cell behaviors^10–14^. Most assays for probing chromatin organization at the molecular level, such as assay for transposase-accessible chromatin sequencing (ATAC-seq), assay for transposase-accessible chromatin with visualization (ATAC-see)^15,16^ or Hi-C^17^, rely on cell fixation or lysis and downstream sequencing. While these methods provide high-resolution measurements of chromatin molecular organization, they cannot assess chromatin reorganization in living cells.

Quantitative live-cell measurements of chromatin compaction have been achieved using sophisticated approaches such as FLIM-FRET using genetically encoded histone reporters^18,19^ or DNA-binding fluorescent dyes^20,21^, and label-free interferometric scattering microscopy^22,23^, but these methods typically require engineered reporter systems, specialized probes, or instrumentation. In contrast, live-cell DNA staining with membrane-permeable dyes^24^ is simple, accessible to all cell types including genetically intractable immune cells, and readily compatible with standard fluorescence microscopy. However, although DNA images are routinely used to visualize chromatin reorganization, fixation-free quantitative readouts derived from such images remain underexplored.

Among simple image-derived descriptors, the coefficient of variation (CV) of nuclear DNA intensity has been used as a proxy for chromatin compaction, including in microscopy-based workflows for quantifying chromatin organization^2,25,26^. CV captures the dispersion of pixel intensities and has been applied as a simple and interpretable descriptor of chromatin organization. However, its performance relative to other image-derived metrics in the context of live-cell chromatin reorganization has not been systematically examined. More broadly, simple metrics derived from nuclear intensity distributions have not been directly benchmarked against one another in a common live-cell framework.

Here, we developed two new metrics to assess chromatin reorganization in live cells—1-Gini and the Diffuse Signal Index (DSI)—and benchmarked them alongside CV using NETosis as a model of chromatin reorganization. NETosis provides a useful testbed because it features a pronounced and molecularly regulated compact-to-decompact transition of chromatin^27–29^. We show that all three metrics track compact-to-decompact chromatin reorganization in live-cell trajectories, while DSI best discriminates NETing from non-NETing cells at the trajectory level. To assess the biological interpretability of CV, DSI, and 1-Gini, we compared them with ATAC-see-based chromatin accessibility measurements in fixed cells and found that all three metrics correlate with chromatin accessibility, supporting their use as fixation-free proxies for chromatin compaction state. To assess the broad applicability of the metrics, we used mitotic U2OS cells and showed that all three metrics capture mitotic chromatin compaction and post-mitotic decompaction. Together, these analyses establish a practical benchmark for fixation-free, image-derived quantification of chromatin organization dynamics in living cells. To facilitate practical implementation, we also provide NucMetrics, an open-source ImageJ/Fiji macro toolset for computing all three metrics without requiring programming.

## Results

### CV, 1-Gini, and DSI quantify chromatin compaction state as complementary measures of intranuclear DNA signal homogeneity

To benchmark simple image-derived metrics for chromatin-state tracking, we compared three metrics derived from nuclear DNA staining: the coefficient of variation (CV), 1-Gini, and the Diffuse Signal Index (DSI) (see Materials and Methods). Briefly, CV is defined as the ratio of the standard deviation to the mean of the raw pixel intensities and captures the overall spread of the intensity histogram. 1-Gini is derived from the Lorenz curve of the min-max normalized pixel distribution and DSI quantifies the fraction of min-max normalized pixels exceeding a threshold of τ. CV decreases with signal uniformity while DSI and 1-Gini, bound between 0 and 1, increase with signal uniformity.

These metrics were selected because they quantify related but non-identical features of intranuclear DNA intensity distribution and are therefore expected to differ in their sensitivity to specific patterns of signal redistribution (see Materials and Methods). To assess how each metric responds across chromatin reorganization states, we compared CV, 1-Gini and DSI in representative nuclei at early and late chromatin reorganization stages during NETosis (**Figure 1**). In the early stage, the DNA signal appeared heterogeneous, with prominent peripheral enrichment (**Figure 1A**), whereas in the late stage it became more uniformly distributed throughout the nuclear ROI (**Figure 1B**). We found that the three metrics capture distinct aspects of the intranuclear intensity distribution (**Figure 1C–H**). CV decreased as chromatin reorganized from the early to the late state (early: CV = 0.583; late: CV = 0.326), reflecting narrowing of the intensity distribution (**Figure 1C, D**). 1-Gini increased with chromatin reorganization (early: 1-Gini = 0.620; late: 1-Gini = 0.797) reflecting greater signal homogeneity as illustrated by the Lorenz curves and the conceptual grid diagram in **Figure 1E** & **F**. DSI with τ = 0.3, increased with chromatin reorganization (early: DSI = 0.367; late: DSI = 0.792), offering the largest dynamic range among the three metrics (**Figure 1G, H**). To select τ, we performed a systematic threshold sweep across paired nuclei at early versus late stages of NETosis and identified τ = 0.3 as the value that maximized discrimination between the two states (**Supplementary Figure 1**).

**Figure 1.**
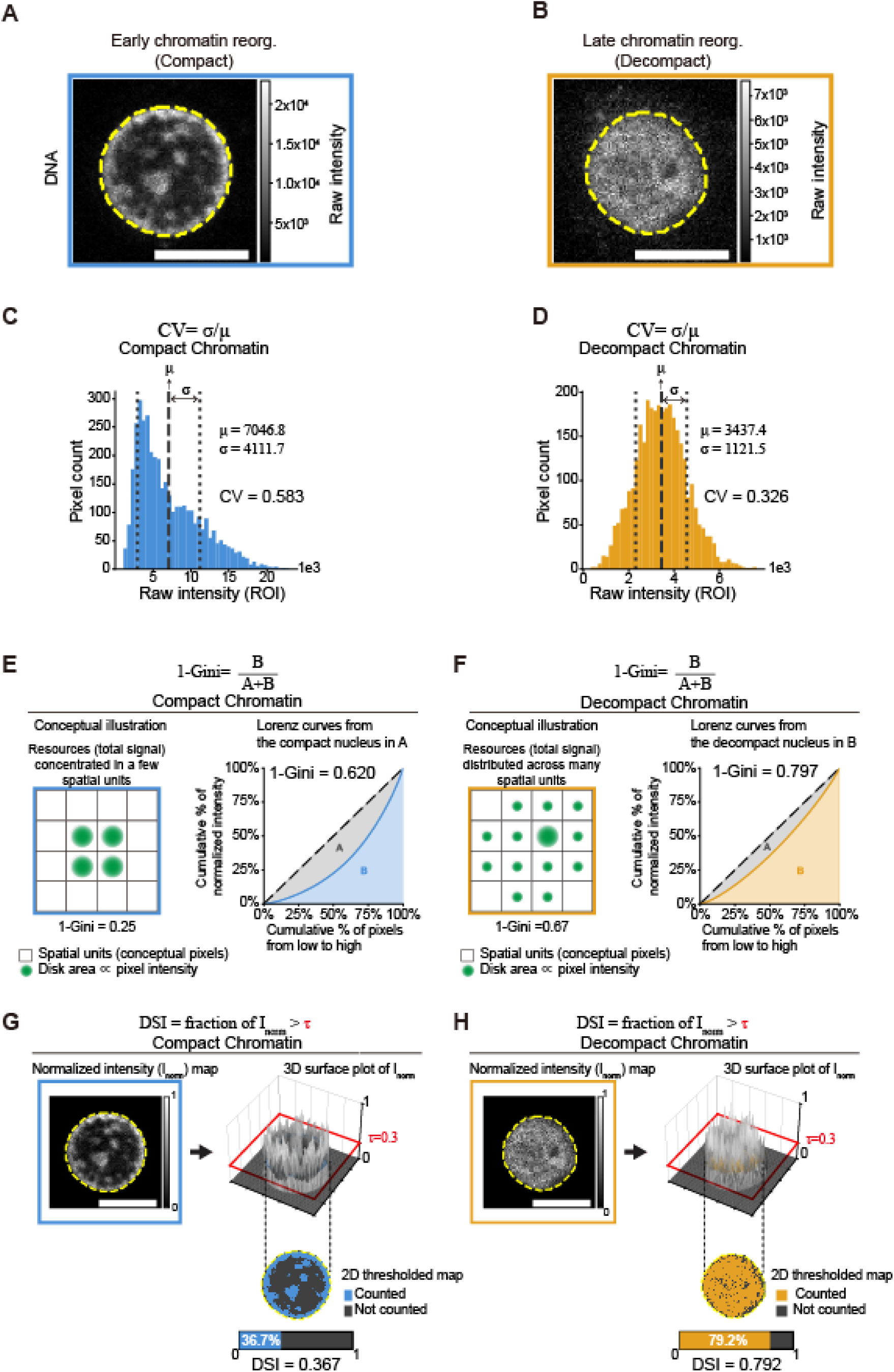
CV, 1-Gini, and DSI quantify chromatin compaction as complementary measures of intranuclear DNA signal homogeneity. **(A, B)** Representative fluorescence images of dHL-60 cell nuclei at early chromatin reorganization (compact; blue box, A) and late chromatin reorganization (decompact; orange box, **B**) stages during NETosis, stained with SPY650-DNA. Raw pixel values are shown with corresponding intensity colorbars. Yellow dashed lines indicate nuclear ROI boundaries. Scale bars, 5 µm. These two chromatin states serve as the basis for metric illustration in panels **C–H**. **(C, D)** CV (coefficient of variation; σ/μ) computed from the raw pixel intensity distribution within the nuclear ROI of the compact **(C)** and decompact **(D)** nuclei in **(A, B)**. Histograms show the distribution of raw pixel intensities for each nucleus; vertical lines indicate the mean (dashed) and ±1 SD (dotted). CV decreases from the compact (CV = 0.583) to the decompact state (CV = 0.326), reflecting narrowing of the intensity distribution. **(E, F)** 1-Gini (= B/(A+B)) computed from the Lorenz curve of the min-max normalized pixel distribution for the compact **(E)** and decompact **(F)** nuclei. Left in each panel: conceptual illustration showing how 1-Gini reflects signal uniformity across spatial units — when equal total signal is concentrated in a few spatial units (1-Gini = 0.25) versus distributed across many units (1-Gini = 0.67), 1-Gini increases proportionally. Disk area is proportional to pixel intensity. Right in each panel: Lorenz curves from the example nuclei in **(A, B)**. The diagonal dashed line represents perfect signal uniformity; area A (gray) represents the region between the diagonal and the Lorenz curve, and area B (condition color) the area below the curve. 1-Gini increases from the compact (1-Gini = 0.620) to the decompact state (1-Gini = 0.797), reflecting greater signal homogeneity.| **(G, H)** DSI (Diffuse Signal Index; fraction of *I_norm_* > τ) computed as the fraction of min-max normalized pixels exceeding a threshold τ = 0.3 for the compact **(G)** and decompact **(H)** nuclei. For each condition, the normalized intensity (*I_norm_*) map is shown alongside a 3D surface plot of *I_norm_* in which the red plane marks τ = 0.3; pixels whose intensity exceeds this plane are counted toward DSI. Below, the 2D thresholded map shows counted pixels (*I_norm_* > τ) in condition color and uncounted pixels (*I_norm_* ≤ τ) in dark gray, with the DSI fraction bar indicating the proportion of counted pixels. DSI increases from the compact (DSI = 0.367) to the decompact state (DSI = 0.792).

Because shot noise can bias intensity-distribution metrics in photon-limited images and nuclei with decompacted chromatin have lower signal, we used two approaches to assess if the measured changes in CV, 1-Gini and DSI was due to the decreased signal of decompacted chromatin. First, we simulated Poisson-noise spanning the low-photon regime described by Longo et al.^30^ (see Material and Methods). We found that although the noise shifted the absolute metric values, compact and decompact states remained separated with the same directionality at every tested noise level (**Supplementary Figure 2**), indicating that the experimental differences are not explained by shot noise alone. Second, we estimated the photon count of our confocal images (see Material and Methods) and found that the photon counts of images with decompacted nuclei exceeded ∼200, thus above the limit of 40 set by Longo et al. Together, these analyses show that CV, 1-Gini, and DSI can discriminate between compact and decompact chromatin in NETing cells, independently of DNA signal noise.

### CV, DSI and 1-Gini track progressive chromatin reorganization in live NETing cells, with DSI best at distinguishing NETing from non-NETing cells

To evaluate whether the three metrics can track progressive chromatin reorganization in living cells, we performed time-lapse fluorescence imaging of dHL-60 cells stimulated for NETosis and compared the trajectories of CV, DSI and 1-Gini between NETing and non-NETing cells (**Figure 2A–D**). Cell trajectories were classified as NETing (n = 42) or non-NETing (n = 28) based on the occurrence of microvesicle shedding, a morphological hallmark of NETosis onset^28^. For population-level comparison, NETing cell trajectories were aligned relative to two specific NETosis events in which chromatin reorganizes inside the nucleus^2,28^: nuclear rounding onset (normalized time = 0) and the last frame preceding nuclear rupture (normalized time = 1). For non-NETing cells, trajectories were normalized relative to the start (normalized time = 0) and end (normalized time = 1) of the movie.

**Figure 2.**
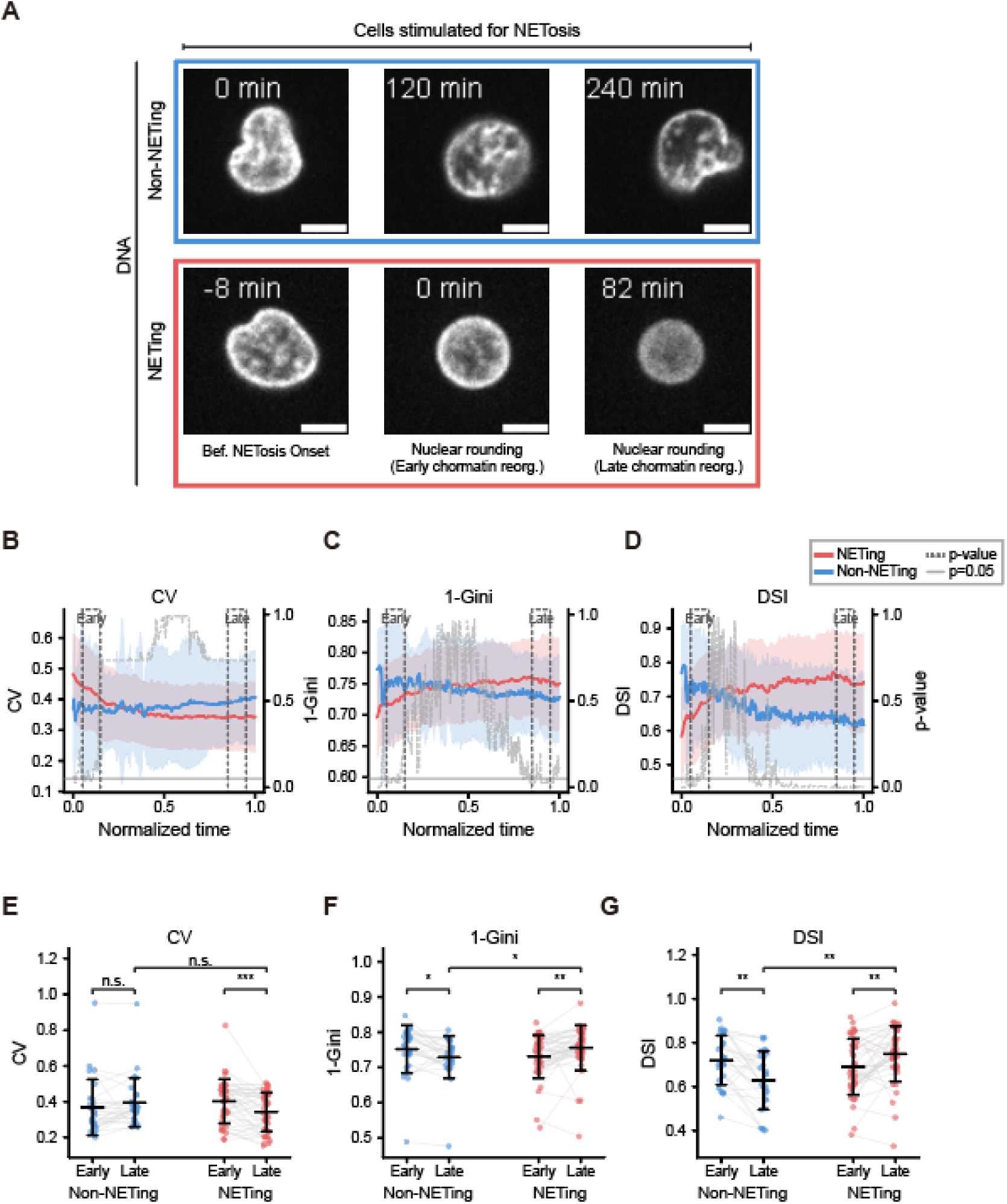
CV, DSI and1-Gini track progressive chromatin reorganization in live NETing cells, with DSI best at distinguishing NETing from non-NETing cells. **(A)** Representative time-lapse fluorescence images of dHL-60 cell nuclei stained with SPY650-DNA during NETosis stimulation (4 µM ionomycin). Top row (blue box): non-NETing cell at 0, 120, and 240 min from movie start. Bottom row (red box): NETing cell at −8, 0, and 82 min relative to nuclear rounding onset. NETosis stages are indicated below each image. Scale bars, 5 µm. **(B–D)** Population-level time series of CV **(B)**, 1-Gini **(C)**, and DSI **(D)** for NETing (red, n = 42) and non-NETing (blue, n = 28) cells. Solid lines indicate population means; shaded regions indicate ±1 SD. Cell trajectories were aligned to normalized time (0 = nuclear rounding onset, 1 = frame immediately preceding nuclear envelope rupture for NETing cells; 0 = movie start, 1 = movie end for non-NETing cells). Gray dashed lines show p-values (right y-axis) at each normalized time point (BH correction); the gray solid line indicates p = 0.05. DSI showed statistically significant separation between NETing and non-NETing trajectories at 62.2% of normalized time points, compared with 17.4% for 1-Gini and 2.9% for CV. **(E–G)** Per-cell early- and late-window means for CV **(E)**, 1-Gini **(F)**, and DSI **(G)** in non-NETing and NETing cells. Each dot represents one cell’s mean metric value within the indicated normalized-time window (early, 0.05–0.15; late, 0.85–0.95), and gray lines connect paired measurements from the same cell. Black horizontal bars and error bars indicate mean ± SD. Brackets show the within-population early-versus-late comparisons and the late-window comparison between NETing and non-NETing cells. Statistical significance was assessed using paired Wilcoxon signed-rank tests for early-versus-late comparisons and a Mann-Whitney U test for the late-window between-population comparison, with Holm correction across the three planned comparisons separately for each metric. All three metrics changed significantly from early to late in NETing cells. At the late window, 1-Gini and DSI, but not CV, differed significantly between populations.

Analysis of CV, DSI and 1-Gini trajectories in NETing cells showed that all three metrics change in manner consistent with DNA signal homogenization and compact-to-decompact chromatin reorganization (**Figure 2B-D**). CV progressively decreases while DSI and 1-Gini progressively increase. To assess if the three metrics can quantitatively track NETosis chromatin reorganization, we calculated and compared their mean per cell values between early and late stages of chromatin reorganization (**Figure 2E-G**, see Material and Methods). We found that CV significantly decreased from 0.427 ± 0.127 to 0.343 ± 0.109; DSI and 1-Gini significantly increased from 0.667 ± 0.131 to 0.750 ± 0.126 and from 0.723 ± 0.063 to 0.756 ± 0.064, respectively. Thus, all three metrics detected the compact-to-decompact transition within NETing cells. Similar analysis of non-NETing cells showed that CV did not significantly change (0.372 ± 0.196 to 0.395 ± 0.136), while DSI and 1-Gini significantly decreased (0.726 ± 0.141 to 0.629 ± 0.133 and 0.751 ± 0.086 to 0.729 ± 0.060, respectively). This decrease in 1-Gini and DSI and the upward trend of CV in non-NETing cells indicate nuclear DNA heterogenization, potentially due to a previously underappreciated early chromatin compaction between NETosis stimulation and NETosis initiation. Together, this data indicate that CV, DSI and 1-Gini can effectively track chromatin decompaction during NETosis.

We next set out to test if our metrics can distinguish populations of NETing from non-NETing cells. We compared the trajectories of CV, DSI and 1-Gini between these two populations at each normalized time point (**Figure 2B-D**). We found that DSI trajectories differed significantly for 62.2% of normalized time points, compared with 17.4% for 1-Gini and 2.9% for CV, with the strongest divergence observed at later time points approaching nuclear rupture. To quantify this divergence, we compared the distributions of per cell means of CV, DSI and 1-Gini between NETing and non-NETing cells in late time windows (**Figure 2E-G**). This showed that NETing cells had significantly higher DSI and 1-Gini than non-NETing cells, whereas CV did not significantly differ between NETing and non-NETing cells Together, this data indicates that CV, DSI and 1-Gini can all track chromatin reorganization during NETosis; that DSI and 1-Gini can distinguish populations of NETing from non-NETing cells with DSI being the best at capturing the differences between these two populations.

### CV, DSI and 1-Gini correlate with ATAC-see-based chromatin accessibility

To assess the biological interpretability of CV, DSI and 1-Gini, we compared them to ATAC-see-based measurements of chromatin molecular accessibility in fixed dHL-60 cells stimulated for NETosis (**Figure 3**). We reasoned that if CV, DSI, and 1-Gini capture biologically meaningful aspects of chromatin organization, they should show measurable relationships with an orthogonal readout of chromatin accessibility.

**Figure 3.**
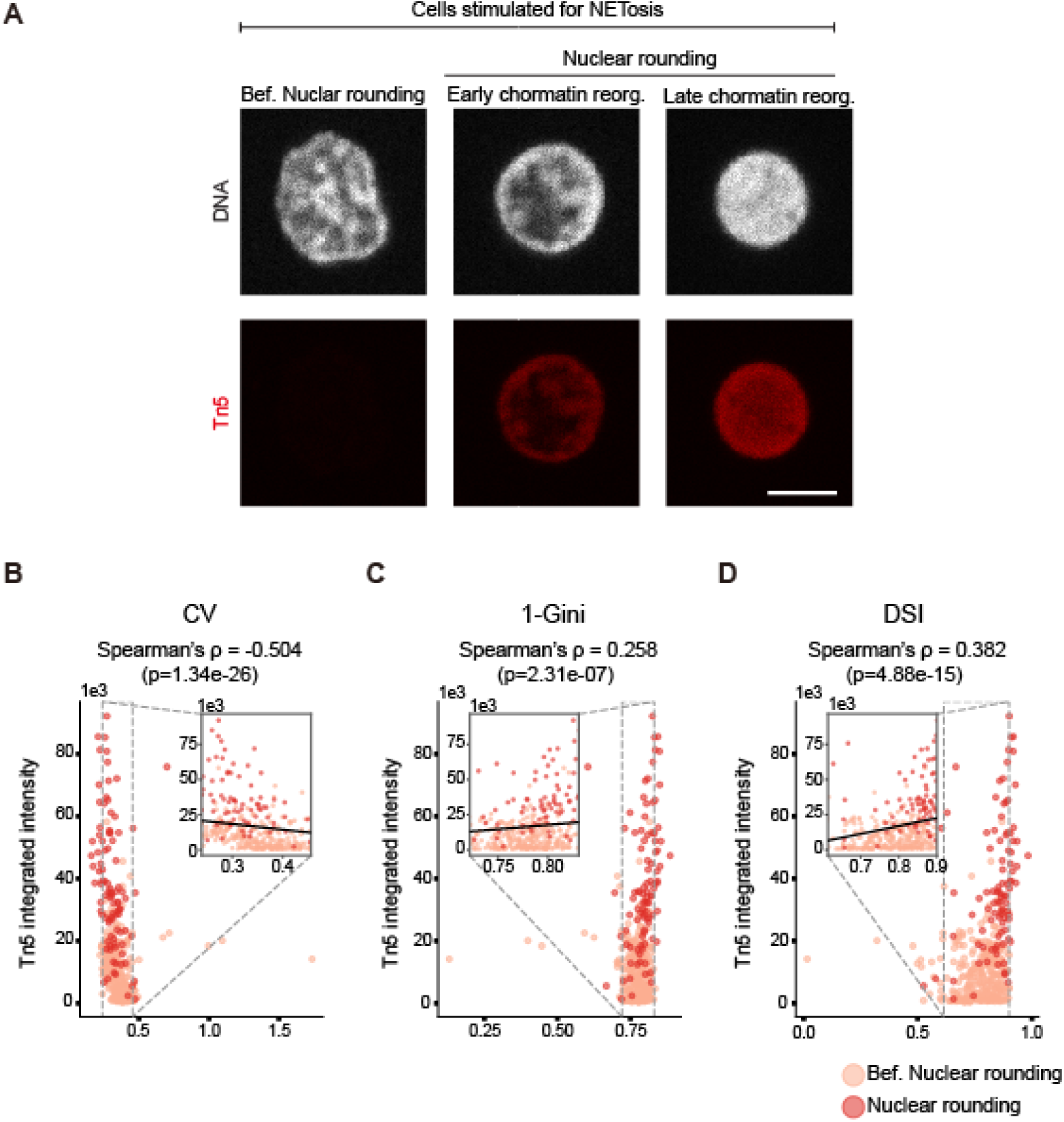
CV, DSI and 1-Gini correlate with ATAC-see-based chromatin accessibility. **(A)** Representative fluorescence images of DNA (DAPI, displayed in grayscale; top) and Tn5 transposase signal (Alexa Fluor 488, displayed in red; bottom) in fixed dHL-60 cells at indicated NETosis stages. NETosis stage was classified based on nuclear morphology and Lamin B Receptor staining (not shown). Scale bar, 5 µm. **(B–D)** Scatter plots of CV **(B)**, 1-Gini **(C)**, and DSI **(D)** versus Tn5 integrated intensity for individual fixed cells at two NETosis stages: before nuclear rounding (light, n = 297) and nuclear rounding (dark, n = 95). Spearman’s ρ and associated p-values are shown above each panel. Insets zoom into the 5th–95th percentile range of each metric on the x-axis (dashed lines indicate the zoomed region) with a linear regression line. All three metrics showed statistically significant monotonic correlations with Tn5-based accessibility. CV exhibited the strongest association (ρ = −0.504), followed by DSI (ρ = 0.382) and 1-Gini (ρ = 0.258); the negative sign of CV reflects its inverse relationship to chromatin decompaction.

Representative fluorescence images showed progressive increase in the ATAC-see signal (Tn5-oligo-AF488) between NETosis-stimulated cells before nuclear rounding and NETing cells at early or late stages of chromatin reorganization **(Figure 3A)**, consistent with increasing chromatin accessibility during NETosis as we previously reported^2^.

We next examined whether CV, DSI and 1-Gini, correlated with Tn5 integrated intensity across a population of NETing cells before and at the nuclear rounding stages of NETosis (n = 297 and 95, respectively; **Figure 3B–D**). All three metrics showed statistically significant Spearman correlations with Tn5 integrated intensity. CV was negatively correlated (ρ = −0.504, p = 1.34 × 10^−26^), consistent with its inverse relationship to DNA signal homogenization, whereas 1-Gini and DSI were positively correlated (ρ = 0.258, p = 2.31 × 10^−7^ and ρ = 0.382, p = 4.88 × 10^−15^, respectively). While the magnitude of these correlations was moderate, the consistent directionality across all three metrics supports their biological interpretability.

Together, these results indicate that CV, DSI and 1-Gini correlate with ATAC-see based chromatin accessibility changes during NETosis; with CV showing the strongest correlation. While these metrics do not directly substitute for molecular accessibility assays, they provide live-cell-compatible image-derived readouts of chromatin organization dynamics.

### CV, DSI and 1-Gini capture bidirectional chromatin reorganization during mitosis is U2OS cells

To test whether CV, DSI and 1-Gini capture chromatin reorganization beyond NETosis, we applied them to U2OS cells undergoing mitosis, a distinct biological process where chromatin compacts at mitotic entry and decompacts at mitotic exit^31^. Because mitotic rounding can transiently displace chromatin from the initial focal plane, we restricted the analysis to manually selected in-focus stages of nuclei in interphase, mitotic entry before nuclear envelope breakdown, and post-mitotic nuclei. We found that all three metrics changed significantly across stages (**Figure 4B–D**). CV increased at mitotic entry from ∼0.208 to ∼0.255 and decreased after mitotic exit to ∼0.173; DSI and 1-Gini decreased at mitotic entry from ∼0.686 to ∼0.557 and from ∼0.806 to ∼0.767, respectively, and increased to ∼0.775 and ∼0.824, respectively in post-mitotic nuclei. These changes are consistent with increased DNA signal heterogeneity during mitotic chromatin compaction followed by signal homogenization during post-mitotic chromatin decompaction. Statistical tests showed that all three metrics changed significantly across the three mitotic stages, with the stage-associated effect size being largest for CV (Kendall’s W = 0.71), followed by 1-Gini (W = 0.53) and DSI (W = 0.44).This suggests that CV is more sensitive to mitosis-related chromatin reorganization than 1-Gini and DSI. Together, these results show that all three metrics capture the expected direction of chromatin reorganization in mitosis, indicating their applicability beyond NETosis.

**Figure 4.**
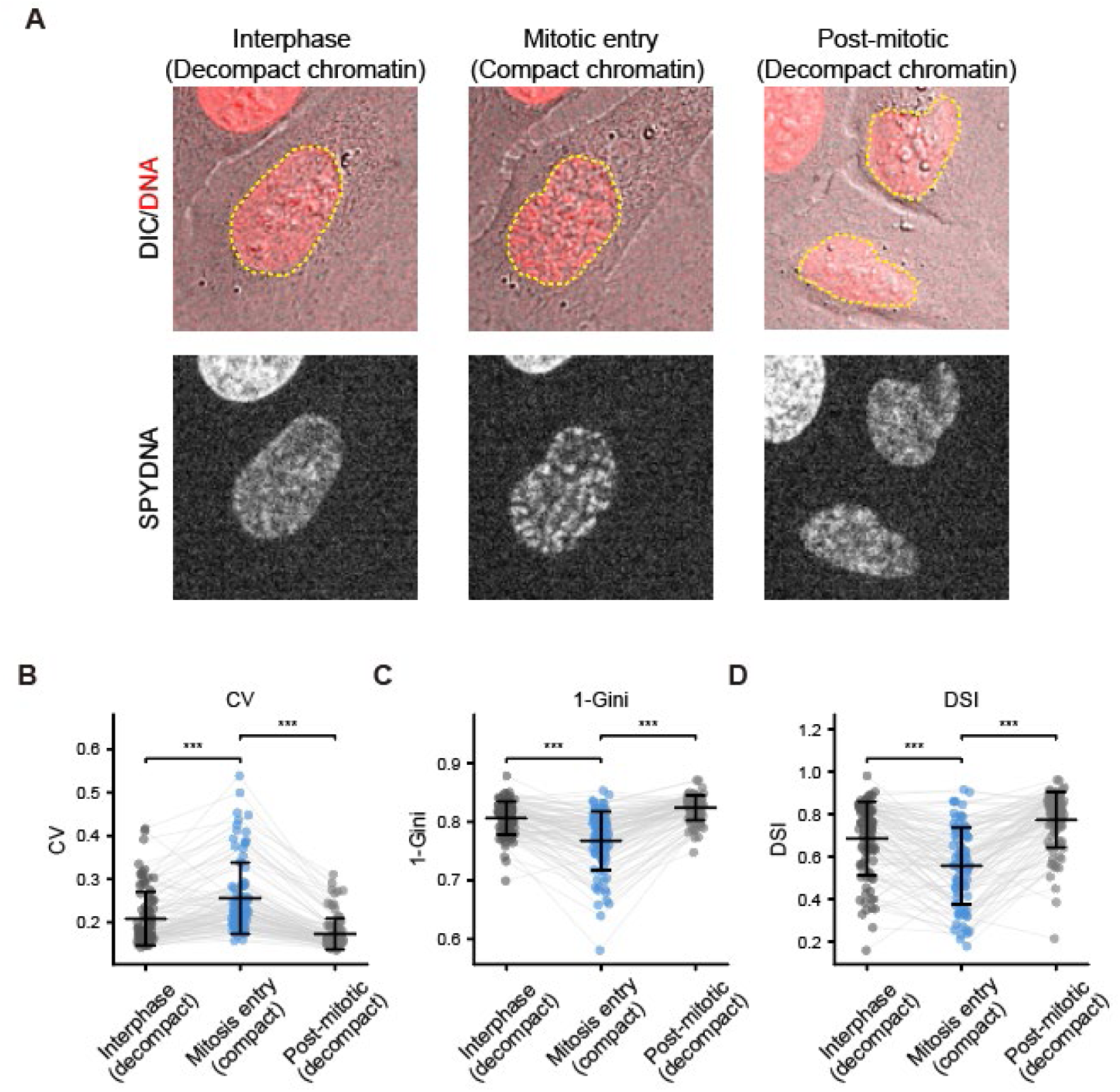
CV, DSI and 1-Gini capture mitotic chromatin compaction and post-mitotic chromatin decompaction in U2OS cells. **(A)** Representative U2OS cell imaged during mitotic progression using live-cell spinning disk confocal microscopy with SPY650-DNA. Images show three manually selected in-focus stages: interphase (decompact chromatin), mitotic entry before nuclear-envelope breakdown (compact chromatin), and post-mitotic daughter nuclei (decompact chromatin). Top row shows merged DIC and SPYDNA fluorescence images; bottom row shows the corresponding SPYDNA channel. Yellow dotted outlines indicate the segmented DNA-containing regions used for metric quantification. Scale bar, 10 µm. **(B–D)** Stage-based quantification of CV **(B)**, 1-Gini **(C)**, and DSI **(D)** across n = 94 cells pooled from three independent experiments. Each dot represents one cell at the indicated stage, and gray lines connect measurements from the same cell across stages. Black horizontal bars and error bars indicate mean ± SD. CV increased from interphase to mitotic entry and decreased after mitotic exit (0.208 ± 0.062, 0.255 ± 0.083, and 0.173 ± 0.036), whereas 1-Gini (0.806 ± 0.028, 0.767 ± 0.050, and 0.824 ± 0.021) and DSI (0.686 ± 0.173, 0.557 ± 0.181, and 0.775 ± 0.131) decreased at mitotic entry and increased in post-mitotic daughter nuclei (interphase, mitotic entry, and post-mitotic; mean ± SD), consistent with chromatin compaction followed by post-mitotic chromatin decompaction. Statistical significance across stages was assessed using a Friedman test (df = 2; CV, χ² = 132.9, p = 1.4 × 10^−29^, Kendall’s W = 0.71; 1-Gini, χ² = 99.4, p = 2.6 × 10^−22^, W = 0.53; DSI, χ² = 82.8, p = 1.0 × 10^−18^, W = 0.44), followed by post hoc paired Wilcoxon signed-rank tests with Holm correction; all pairwise stage comparisons were significant for each metric (all p_adj < 0.05). Kendall’s W is the omnibus effect size for the Friedman test and summarizes the overall stage-associated difference across the three repeated measurements. Brackets in **(B–D)** show adjacent-stage comparisons.

### NucMetrics: an ImageJ/Fiji toolset for accessible computation of CV, 1-Gini, and DSI

To make the metrics described above accessible to a broad audience, we developed NucMetrics, an open-source ImageJ/Fiji macro toolset that computes CV, 1-Gini, and DSI directly within the Fiji environment (**Figure 5**). NucMetrics is distributed as a single macro file (.ijm; see Code Availability) that installs into the Fiji toolbar and requires no external dependencies or plugins (**Figure 5A–C**). The toolset supports four modes of nuclear region definition: (1) manual ROI selection, in which the user draws a freehand or polygon ROI around a nucleus; (2) ROI Manager mode, which batch-processes multiple pre-defined ROIs; (3) binary mask mode, in which the user supplies a pre-existing binary mask image to define nuclear regions; and (4) auto-generate binary mask mode, which generates a binary mask via automatic thresholding (Li, Otsu, or Triangle), morphological cleanup, and particle analysis, displays the mask for visual review, and computes metrics upon user confirmation (**Figure 5A, C**). Both binary mask and auto-generate modes support whole-stack processing for time-lapse movies of single nuclei, enabling trajectory-level quantification of CV, 1-Gini, and DSI over time. A step-by-step tutorial and example time-lapse dataset are provided in the GitHub repository. For all modes, the three metrics are computed from pixel intensities within the defined nuclear regions using the same formulas described in Materials and Methods. Results, including CV, 1-Gini, DSI, mean intensity, standard deviation, and pixel count for each nucleus, are output to the ImageJ Results table for straightforward export and downstream analysis (**Figure 5F**).

**Figure 5.**
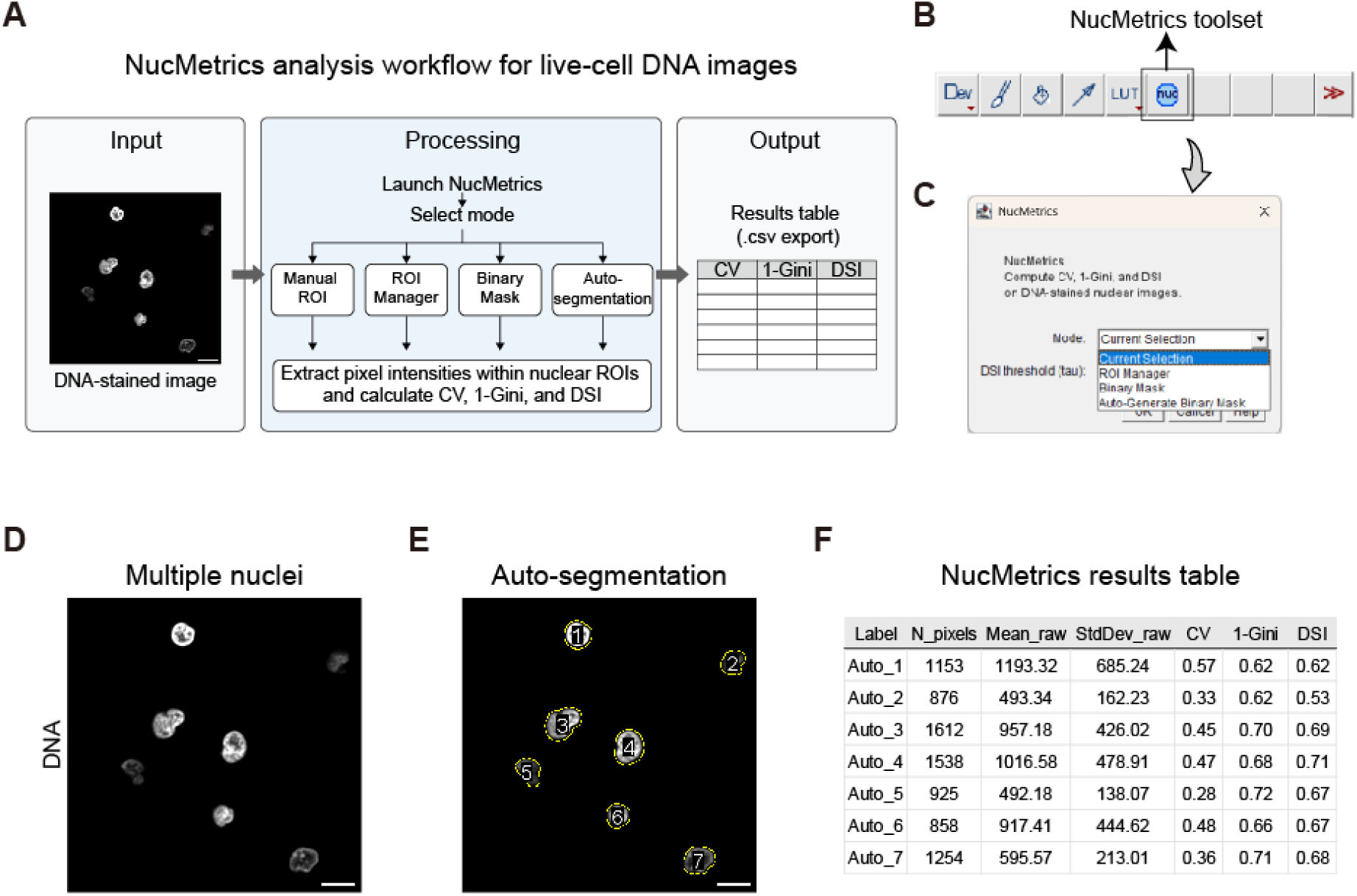
NucMetrics: an ImageJ/Fiji macro toolset for accessible computation of CV, 1-Gini, and DSI. **(A)** Schematic workflow of NucMetrics. The toolset accepts a DNA-stained nuclear image (left) and provides four modes for defining nuclear regions: manual ROI selection, ROI Manager batch processing, external binary mask, and auto-generate binary mask (Li, Otsu, or Triangle thresholding with user review). For each detected nucleus, the macro extracts pixel intensities within the defined region and computes CV (from raw intensities), 1-Gini, and DSI (from min-max normalized intensities, with user-adjustable threshold tau). Results are output to a Results table. **(B)** NucMetrics installed in the Fiji toolbar (arrow). **(C)** The NucMetrics mode selection dialog showing the four available modes (Current Selection, ROI Manager, Binary Mask, Auto-Generate Binary Mask) and the adjustable DSI threshold parameter (tau). A Help button links to the associated publication. **(D–F)** Example application of NucMetrics in auto-generate binary mask mode on a single time point from a live-cell time-lapse of dHL-60 cells stained with SPY650-DNA. **(D)** Original fluorescence image of multiple nuclei. **(E)** The same image with NucMetrics-detected nuclear ROIs overlaid (white outlines; 7 nuclei detected). **(F)** The corresponding Results table showing CV, 1-Gini, and DSI computed for each nucleus. Scale bars, 10 µm.

As an example of how NucMetrics is used in Fiji, and of the outputs it generates, we applied NucMetrics in auto-generate binary mask mode to a single time point from a live-cell time-lapse movie of dHL-60 cells stained with SPY650-DNA (**Figure 5D–F**). NucMetrics automatically identified 7 nuclei and displayed the detected ROIs overlaid on the original image for visual verification (**Figure 5E**). The corresponding Results table shows CV, 1-Gini, and DSI computed for each detected nucleus (**Figure 5F**). Thus, NucMetrics allows easy calculation of CV, 1-Gini, and DSI for simple assessment of chromatin organization in live cells.

## Discussion

Here, we benchmarked three simple image-derived metrics—CV, DSI, and 1-Gini—for fixation-free quantification of chromatin organization from live-cell DNA images, using NETosis as a model system with a pronounced compact-to-decompact transition. Our results show that all three metrics track compact-to-decompact chromatin reorganization in NETing cell trajectories and correlate with ATAC-see based measurements of chromatin accessibility while differing in their ability to discriminate NETing from non-NETing cells. We further show that the metrics capture mitotic chromatin compaction and post-mitotic chromatin decompaction in U2OS cells, extending the benchmark to a second biological context. Thus, even simple metrics derived from routine DNA staining images can provide informative readouts of chromatin organization in living cells.

A key finding of this study is that all three simple image-derived metrics—CV, DSI and 1-Gini—can track chromatin decompaction during NETosis as well as chromatin compaction then decompaction in mitotic cells. Paired early-to-late NETosis chromatin reorganization analysis showed significant changes in all three metrics within individual NETing cells. When late-window values were compared between NETing and non-NETing cells, DSI showed the strongest separation, followed by 1-Gini, whereas CV did not significantly distinguish between the two populations. This relative ranking was, however, specific to the NETosis benchmark. In mitotic U2OS cells, the overall stage-associated effect was largest for CV, followed by 1-Gini and DSI (**Figure 4**). This different ranking supports the view that CV, DSI and 1-Gini provide complementary readouts whose relative response magnitude depends on the biological context and analytical task. The better performance of DSI in NETosis may reflect its threshold-based design. By quantifying the fraction of pixels above a defined normalized intensity threshold, DSI assesses the expansion of diffuse signal during chromatin reorganization and may therefore be particularly sensitive to the compact-to-decompact transition captured in live-cell NETosis trajectories. Moreover, because the threshold τ can be adjusted to suit different cell types, chromatin perturbations, or imaging conditions, DSI offers a tunable framework that users can optimize for their specific experimental needs.

All three metrics showed statistically significant correlations with ATAC-see signal, with CV showing the strongest association with ATAC see-based chromatin accessibility (ρ = −0.504), followed by DSI (ρ = 0.382) and 1-Gini (ρ = 0.258). This demonstrates that CV, 1-Gini and DSI can capture changes in chromatin accessibility. The moderate correlation might be due to the reduced sensitivity of these metrics in fixed cells. Alternatively, CV, DSI and 1-Gini might not have the sensitivity necessary to capture molecular accessibility. Indeed, CV, DSI and 1-Gini, quantify the mesoscale distribution of DNA fluorescence signal across the nucleus. In contrast, ATAC-see reports molecular-scale chromatin accessibility - the ability of fluorescently labelled Tn5-oligo to bind accessible chromatin regions. Because mesoscale DNA signal redistribution and molecular-scale chromatin accessibility operate at different spatial resolutions, perfect correspondence might not be achievable. Nevertheless, the consistent directionality of the correlations between the ATAC-see signal and the three metrics supports that these metrics capture image features meaningfully linked to the state of chromatin accessibility.

Several limitations should be noted. First, this benchmarking was performed in two biological systems, NETosis in dHL60 and mitosis in U2OS cells, chosen because they provide a large and physiologically relevant chromatin transition. Future work is needed to establish universal metric performance in other contexts, including subtler chromatin remodeling programs. Second, the live image-based analyses were performed in 2D, and future work will be needed to assess how these metrics behave in volumetric imaging datasets. Third, the DSI threshold was optimized for the present system and imaging conditions, and may require recalibration for different cell types, chromatin perturbations, DNA dyes, or microscopy platforms. Fourth, intensity-based metrics are sensitive to shot noise^30^ and although our simulation indicates that the CV, DSI and 1-Gini can still distinguish compact from decompacted chromatin in presence of noise, imaging conditions should be tuned to optimize signal to noise ratio. Fifth, all three metrics operate at the mesoscale—quantifying bulk redistribution of DNA fluorescence signal within the nucleus—and do not resolve molecular-scale chromatin features such as nucleosome positioning. They therefore complement, rather than replace, molecular-resolution approaches such as ATAC-seq, ATAC-see, or FLIM-FRET.

Despite these limitations, the practical value of this framework lies in its simplicity and accessibility. CV, 1-Gini, and DSI can all be extracted from standard live-cell DNA imaging without fixation, engineered reporters, or specialized instrumentation. As such, they provide an experimentally straightforward bridge between qualitative visualization of chromatin reorganization and quantitative analysis of chromatin dynamics. To ensure that our metrics are accessible to a broad audience, we developed NucMetrics, an open-source ImageJ/Fiji macro toolset that computes all three metrics without requiring programming experience. Thus, this study establishes a simple, accessible and quantitative framework for assessing global chromatin reorganization in living cells.

## Materials and Methods

### Cell culture and differentiation

Human promyelocytic leukemia HL-60 cells (ATCC, CCL-240) were maintained in RPMI 1640 medium (Cytiva, SH30096.01) supplemented with 1% GlutaMAX (Thermo Scientific, 35050061), 1% Penicillin/Streptomycin (Gibco, 15070063), 25mM HEPES (Cytiva, SH30237.01), and 15% heat-inactivated fetal bovine serum (FBS; R&D Systems, S11150). Cells were passaged every 3 days at 2 × 10^5^ cells/mL and maintained at 37°C in a humidified 5% CO_2_ incubator. To differentiate HL-60 cells into neutrophil-like dHL-60 cells, culture medium was supplemented with 1.3% dimethyl sulfoxide (DMSO; Sigma-Aldrich, D2650). Differentiated cells were used for experiments on day 6 or day 7 post-DMSO treatment.

Human osteosarcoma U2OS cells (ATCC, HTB-96) were cultured in McCoy’s 5A medium (Cytiva, SH30200.01) supplemented with 1% Penicillin/Streptomycin (Gibco, 15070063) and 10% fetal bovine serum (FBS; R&D Systems, S11150). Cells were passaged every 3 days or at 80–90% confluency using 0.25% trypsin-EDTA (Thermo Scientific, 25200072), seeded at 2 × 10^5^ cells in tissue-culture-treated 25 cm^2^ flasks (CELLTREAT, 229331), and maintained at 37°C in a humidified 5% CO_2_ incubator.

### dHL-60 live-cell sample preparation and NETosis induction

For live-cell imaging, dHL-60 cells (day 6 or 7 post-differentiation) were stained with 1µM SPY650-DNA (Cytoskeleton, CY-SC501; reconstituted in DMSO) for 1 hour at 37°C in a humidified 5% CO_2_ incubator. Cells were then pelleted at 2000 rpm for 5 minutes at room temperature and resuspended in imaging medium consisting of serum-free, phenol red-free RPMI 1640 (Gibco, 11835030) supplemented with 25 mM HEPES, 1% Penicillin/Streptomycin, and 1× GlutaMAX. Approximately 105 cells were plated on a non-coated, gamma-irradiated glass-bottom 35 mm dish (World Precision Instruments, FD35-100) or 4-well chambered coverglass (CellVis, C4-1.5HN) pre-equilibrated on a heated microscope stage (37°C). Cells were allowed to adhere for 5 minutes before image acquisition was initiated. After 3–5 minutes of baseline imaging, ionomycin calcium salt (Sigma-Aldrich, I0634; 4µM final concentration) was added to stimulate NETosis and imaging was resumed.

### U2OS live-cell sample preparation

For live imaging of mitotic progression, U2OS cells were stained with 1 µM SPY650-DNA (Cytoskeleton, CY-SC501) together with 10 µM verapamil (Spirochrome; a P-glycoprotein efflux-pump inhibitor used to enhance intracellular SPY650-DNA retention) in complete McCoy’s 5A medium for 1 hour at 37°C in a humidified 5% CO_2_ incubator. Cells were then washed and switched to fresh complete McCoy’s 5A medium for imaging.

### Fixed-cell sample preparation, immunofluorescence, and ATAC-see labeling

For fixed-cell experiments, approximately 5 × 10^4^ dHL-60 cells in 20 µL of imaging medium were seeded on plasma-cleaned 12 mm diameter #1.5 glass coverslips (EMS, 72290-04) and allowed to adhere for 5–10 minutes at 37°C in a humidified 5% CO_2_ incubator. For NETosis-stimulated samples, 200 µL of imaging medium containing ionomycin calcium salt (Sigma-Aldrich, I0634; 4µM final concentration) was added and cells were incubated for 1 hour at 37°C. Unstimulated cells were fixed immediately after the initial adhesion period. Cells were fixed with 4% paraformaldehyde (PFA; Electron Microscopy Sciences, 15710) in 1× cytoskeleton buffer (CB; 10 mM MES, 138 mM KCl, 3 mM MgCl_2_, 2 mM EGTA) for 20 minutes at 37°C. Unreacted PFA was quenched with 0.1 M glycine in 1× CB, followed by permeabilization with 0.5% Triton-X100 in 1× CB for 10 minutes at 37°C. Cells were washed twice with washing buffer (0.1% Tween-20 in 1× TBS; 20 mM Tris-HCl pH 7.6, 150 mM NaCl) for 5 minutes each at room temperature.

For ATAC-see labeling, permeabilized cells were incubated in tagmentation solution (100 nM Alexa488-Tn5 in 1× TD buffer; 2× TD buffer: 20 mM Tris-HCl pH 7.6, 10 mM MgCl_2_, 20% N,N-dimethylformamide diluted in sterile dH₂O) overnight at 4°C. Cells were then washed three times with ATAC-see washing buffer (0.01% SDS, 50 mM EDTA in 1× DPBS) for 15 minutes each at 55°C. Following Tn5 labeling, cells were immunostained for the nuclear envelope marker Lamin B Receptor (LBR) and counterstained with DAPI to enable NETosis stage classification. Fixed and permeabilized cells were blocked in blocking solution (2% BSA, 0.1% Tween-20 in 1× TBS) for 1 hour at room temperature, then incubated with mouse anti-LBR primary antibody (Abcam, BBmLBR 12.F8; 1:500 in blocking solution) for 2 hours at room temperature. After three washes (5 minutes each) in washing buffer, cells were incubated with Alexa Fluor 594 donkey anti-mouse secondary antibody (Jackson ImmunoResearch, 715-587-003; 1:500) and DAPI (1µg/mL; Sigma-Aldrich, D9542) in blocking solution for 1 hour at room temperature. Cells were washed three times in washing buffer and mounted on glass slides using fluorescence mounting medium (Dako, S302380-2).

The Alexa Fluor 488-conjugated Tn5 transposase was kindly provided by Xie Liangqi (Lerner Research Institute, Cleveland Clinic, Ohio, USA) and assembled as previously described ^2^.

## Microscopy

### Spinning disk confocal microscopy

Live-cell time-lapse imaging and fixed-cell imaging were performed on a Nikon Eclipse Ti2 inverted microscope equipped with Perfect Focus, an Okolab stage-top incubator for controlled temperature, humidity and CO2, a Crest X-Light V3 spinning disk scanhead, a Kinetix sCMOS camera, and a Plan Apo 60× oil 1.4 NA DIC objective lens. Illumination was provided by a 7-line solid-state Celesta laser unit (405, 445, 488, 515, 561, 640, and 730 nm). Microscope functions were controlled with NIS-Elements software (Nikon). For dHL-60 live-cell NETosis imaging, cells were imaged at two focal planes: at the coverslip–cell interface (Z = 0 µm) and 3 µm above (Z = +3 µm). Time-lapse images were acquired every 1 minute for 4 hours. For U2OS live-cell imaging of mitotic progression, cells were imaged in a single focal plane using the 640 nm laser at 10% power. Time-lapse images were acquired every 2 minutes for 16 hours. For fixed-cell imaging of ATAC-see samples, 3D Z-stacks were acquired at 0.3 µm step size covering the entire cell.

### Laser scanning microscopy

Representative static images of nuclei at early and late chromatin reorganization stages (**Figure 1A**) were acquired on a Zeiss LSM 900 Airyscan laser scanning microscope equipped with a Plan-Apochromat 63×, 1.40 NA oil immersion objective. The pinhole was set to 1 Airy unit (56 µm) and illumination was provided using a 640 nm laser at 2% power. Microscope functions were controlled with ZEN software (Zeiss).

### Image processing and nuclear segmentation

All image analysis was performed using custom Python scripts (NumPy, SciPy, scikit-image, OpenCV, tifffile, pandas, and matplotlib). Nuclear segmentation was performed on the DNA channel (SPY650-DNA or DAPI) using histogram mode-based intensity thresholding. Briefly, for each image, a threshold was defined as the histogram mode intensity plus a factor of k·SD (k = 1.0–2.0, empirically optimized on pilot datasets and held constant thereafter). Objects smaller than 100 pixels were excluded. For fixed-cell images acquired as 3D Z-stacks, the analysis focal plane (Z_selected) was determined as the slice with the maximum summed nuclear area across the stack. Binary nuclear masks were generated from Z_selected using the same mode + k·SD thresholding, followed by removal of small objects and binary hole filling. For live-cell time-lapse images, nuclear masks were generated at each time point using the same thresholding approach applied to the DNA channel. All nuclear masks were visually inspected and manually corrected where necessary. For dHL-60 live-cell and fixed-cell analyses, downstream metric calculations were performed exclusively on pixels within the nuclear mask.

### U2OS mitotic stage analysis

For U2OS mitotic progression analysis, CV, 1-Gini, and DSI were computed within manually segmented DNA-containing regions at three manually selected in-focus stages: interphase (decompact chromatin), mitotic entry before nuclear-envelope breakdown (compact chromatin), and post-mitotic daughter nuclei (decompact chromatin). Differences across stages were assessed using a Friedman test followed by post hoc paired Wilcoxon signed-rank tests with Holm correction.

### Calculation of CV, 1-Gini and DSI

The three metrics were computed from the pixel intensity distribution within the nuclear mask. Let *I_i_* denote the raw intensity of the pixel *i* within the nuclear mask, where *i* = 1,…,*N* and *N* is the total number of masked pixels.

For 1-Gini and DSI, raw masked pixel intensities were first min-max normalized to the range [0, 1] as

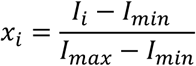

where *I_min_* and *I_max_* are the minimum and maximum raw pixel intensities within the nuclear mask, respectively. The resulting normalized values *xi*∈[0,1] were used for 1-Gini and DSI calculations. CV was computed directly from the raw masked pixel intensities without normalization.

Coefficient of variation (CV) was calculated as

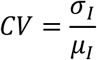

where *σ_I_* and *μ_I_* are the standard deviation and mean of the raw masked pixel intensities, respectively.

The Gini coefficient, originally developed in economics ^32^, quantifies inequality in a distribution. The Gini coefficient was calculated from the normalized pixel intensity distribution as

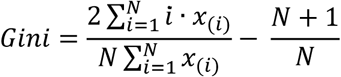

where *x*_(*i*)_ are the normalized pixel intensities sorted in ascending order. The reported metric is 1−Gini, bounded between 0 and 1, where higher values indicate more uniform signal distribution.

The Diffuse Signal Index (DSI) was computed from the min-max normalized pixel distribution as the fraction of masked pixels exceeding a threshold τ:

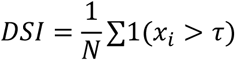

where 1(⋅) is the indicator function, *N* is the total number of masked pixels, and τ is the threshold applied to the normalized intensity distribution. DSI ranges from 0 to 1, with higher values indicating a greater fraction of nuclear pixels occupied by diffuse high-intensity signal and thus a more spatially homogenized intranuclear DNA pattern. Unless otherwise indicated, τ = 0.3 was used. The selection of τ = 0.3 is described in **Supplementary Figure 1**.

### Time normalization and trajectory classification

Cell trajectories were classified as NETing or non-NETing based on the occurrence of microvesicle shedding following ionomycin stimulation. Cells that exhibited microvesicle shedding were classified as NETing, as this event marks the onset of NETosis^28^. Cells that showed no microvesicle shedding throughout the imaging period were classified as non-NETing. For metric extraction, NETing cell trajectories were defined from nuclear rounding onset to the frame immediately preceding nuclear rupture, at which point nuclear tracking was discontinued. Non-NETing cell trajectories spanned the full duration of the movie.

To enable population-level comparison across asynchronous cells, each trajectory was mapped onto a normalized time axis ranging from 0 to 1. For NETing cells, normalized time 0 was defined as the onset of nuclear rounding and normalized time 1 as the frame immediately preceding nuclear rupture. For non-NETing cells, normalized time 0 and 1 corresponded to the first and last frames of the movie, respectively. Normalized time was computed as

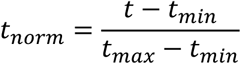

where *t_min_* and *t_max_* are the first and last time points of each cell’s trajectory as defined above.

Two distinct analyses were performed on the live-cell trajectories. For the population-level time-series analysis (**Figure 2B–D**), each cell trajectory was linearly interpolated onto a common 241-point normalized time grid, and the population mean ± SD was computed at each time point. For the per-cell early/late analysis (**Figure 2E–G**), each interpolated cell trajectory was summarized by two values: the mean metric value over an early normalized-time window (0.05–0.15) and the mean over a late window (0.85–0.95). A window value was retained when at least three finite interpolated values were available. Early and late values were paired within each cell, and group summaries are reported as mean ± SD across cells.

### Signal-level estimate and shot-noise simulation

To assess whether the representative spinning disk confocal images used in this study were acquired in a low-photon-count regime, we estimated detected photoelectrons in our raw confocal images. Images were acquired using a Kinetix sCMOS camera in 16-bit Dynamic Range mode with a 2×2 binning. Because images were not acquired with a photon-counting detector, raw pixel intensities were treated as analog-to-digital units (ADU), not direct photon counts. For each representative image, the median intensity of the non-nuclear background region was subtracted from each nuclear pixel, and the resulting background-corrected ADU values were converted to detected photoelectrons using the manufacturer-reported Dynamic Range conversion gain of 0.23 electrons/ADU:

Estimated detected electrons = (ADU_nuclear − ADU_background median) × 0.23.

Quantum efficiency correction was not applied; therefore, the estimated detected-electron values represent a conservative lower-bound proxy for photons reaching the camera sensor. Because the analyzed images were stored after 2×2 binning, the primary estimate is reported per stored 2×2-binned image pixel. As an additional conservative estimate, detected-electron values were also divided by four to approximate the signal per physical sensor pixel.

To test how photon-count-dependent shot noise affects CV, 1-Gini, and DSI, we performed controlled Poisson shot-noise simulations on matched compact and decompact nuclear images from the same cell (**Supplementary Figure 2**). The original nuclear masks were kept fixed for all simulated images so that the analysis tested metric sensitivity, not segmentation robustness. For each image, raw intensities were min-max normalized, scaled to a defined maximum photon budget, and Poisson-sampled. Specifically, the brightest pixel was assigned either 200 or 40 expected photons, and all other pixels were scaled proportionally. The Poisson-sampled image was then rescaled back to the original intensity range before calculating CV, 1-Gini, and DSI using the same formulas as for the experimental data. Photon shot noise was modeled as Poisson noise only; sCMOS read noise was not added. For each photon-budget condition, 500 independent simulations were performed using a fixed random seed. Metric values are reported as mean ± SD across simulations.

### NucMetrics ImageJ/Fiji toolset

NucMetrics was implemented as a single ImageJ macro language (IJM) file by reimplementing the metric formulas from our Python analysis pipeline into the ImageJ macro language. The core metric calculations—CV from raw pixel intensities, 1-Gini and DSI from min-max normalized pixel intensities—use the same mathematical formulas described above. Nuclear segmentation in auto-generate mode uses ImageJ’s built-in functions: automatic intensity thresholding (Li, Otsu, or Triangle), binary hole filling, morphological opening, and particle analysis. Implementation accuracy was verified by comparing CV, 1-Gini, and DSI outputs from NucMetrics against those from our Python analysis code on the same test images. CV outputs were additionally cross-checked by an independent user (A.T.C.) against previously computed CV values from separate NETosis datasets. NucMetrics requires Fiji (ImageJ 1.53b or later) with no additional plugins or dependencies (see Code Availability). Both binary mask and auto-generate modes support whole-stack processing for time-lapse movies of single nuclei; an example time-lapse dataset and step-by-step trajectory quantification tutorial are provided in the GitHub repository.

### Statistical analysis

All statistical analyses were performed in Python using SciPy (scipy.stats). For time-series comparison of NETing and non-NETing trajectories, a Brunner-Munzel test was performed independently at each normalized time point and the resulting p-values were corrected using the Benjamini-Hochberg false discovery rate procedure (**Figure 2B–D**). For the per-cell early/late analysis (**Figure 2E–G**), early and late values were compared within non-NETing and NETing cells using two-sided paired Wilcoxon signed-rank tests, and late-window values were compared between NETing and non-NETing cells using a two-sided Mann-Whitney U test. The three planned comparisons were corrected by the Holm method separately for each metric. Correlations between image-derived metrics and Tn5 integrated intensity were assessed using Spearman rank correlation (**Figure 3B–D**). For U2OS mitotic progression, paired stage-based comparisons across interphase, mitotic entry, and post-mitotic daughter nuclei were assessed using a Friedman test. Kendall’s W was calculated as χ²/[n(k−1)] and reported as the omnibus effect size; W ranges from 0 to 1 and summarizes the consistency of the overall stage-associated difference across cells, but does not indicate the direction of change or identify a specific pairwise contrast. Post hoc paired Wilcoxon signed-rank tests with Holm correction were used for pairwise stage comparisons (**Figure 4B–D**). For DSI threshold selection, paired compact and decompact nuclei were compared using the Wilcoxon signed-rank test at each tested threshold (**Supplementary Figure 1**). P-values less than 0.05 were considered statistically significant. Multiple-testing correction was applied as described above.

## Availability of data and materials

All data used to generate the figures in this study are publicly available through the BioImage Archive under accession number S-BIAD3245 (https://doi.org/10.6019/S-BIAD3245).

## Code availability

All analysis code, the NucMetrics ImageJ/Fiji macro toolset, and example data used for toolset validation are freely available on this GitHub link.

## Acknowledgments

We thank members of the Thiam lab for constructive feedback. We thank Dr. Shannon Yan for helpful discussions.

## Author contributions

M.K. and H.R.T. conceived the research and wrote the manuscript. M.K. conducted the study, and prepared figures. A.T.C. implemented the CV metric for NETosis. M.S. performed ATAC-see experiments. H.R.T. supervised the study.

---

**Supplementary Figure 1.**
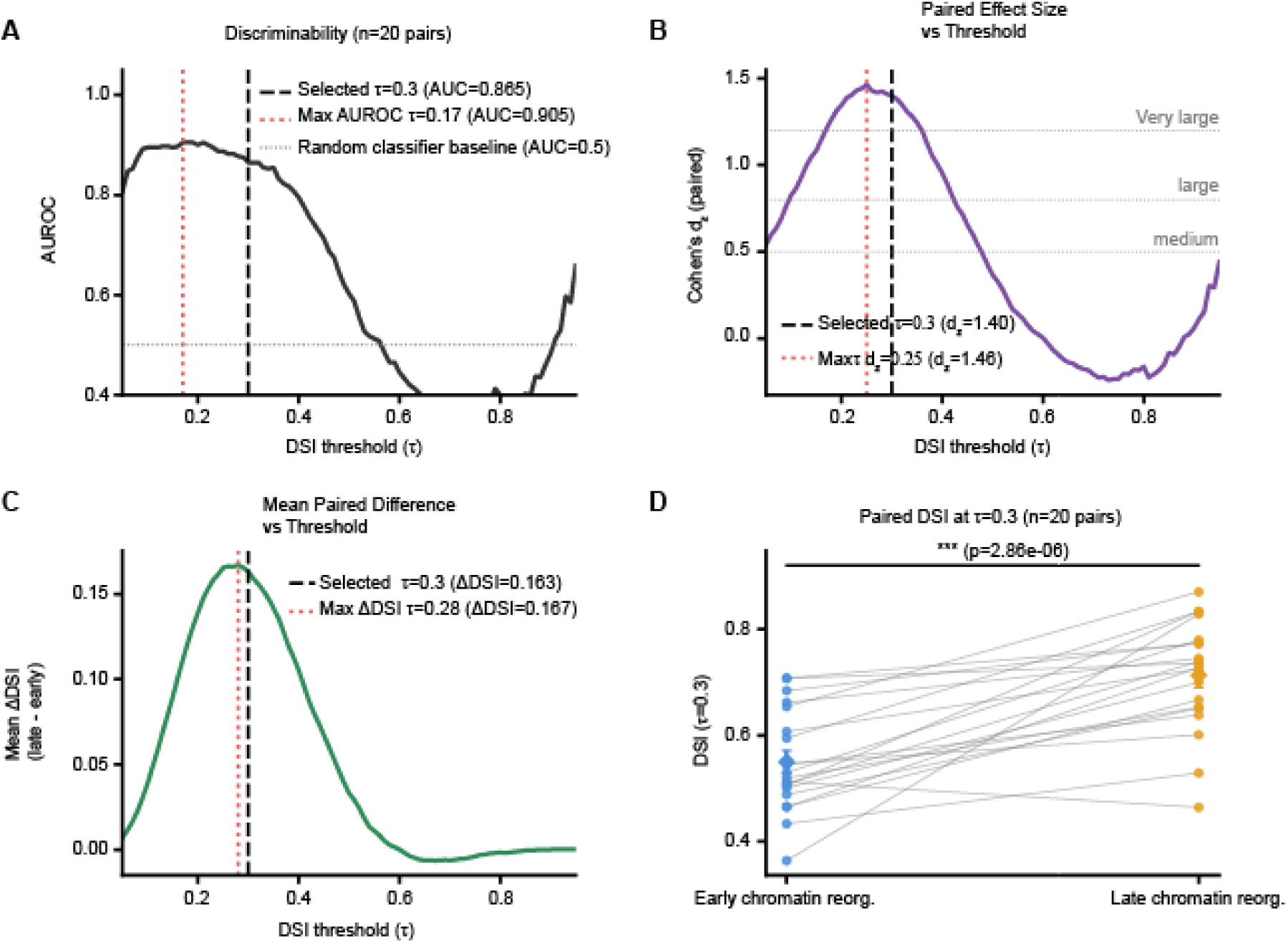
Systematic threshold sweep for DSI threshold selection in paired dHL-60 nuclei. **(A)** DSI threshold sensitivity analysis across 20 paired dHL-60 nuclei imaged at early chromatin reorganization (compact) and late chromatin reorganization (decompact) states from the same nucleus. For each threshold *τ*ranging from 0.05 to 0.95, DSI was computed after strict ROI-wise min-max normalization of DNA intensity. AUROC is plotted as a function of *τ*. The dashed black line indicates the selected threshold (*τ* = 0.3), the dotted red line indicates the threshold yielding the maximum AUROC (*τ* = 0.17), and the gray dotted horizontal line marks the random-classifier baseline (AUROC = 0.5). AUROC at the selected threshold was 0.865. **(B)** Cohen’s *d_z_* for paired differences in DSI between late and early chromatin reorganization states, plotted as a function of threshold *τ*. The dashed black line indicates the selected threshold (*τ* = 0.3), and the dotted red line indicates the threshold yielding the maximum paired effect size (*τ* = 0.25). Gray dotted horizontal lines mark conventional reference levels for medium, large, and very large effect sizes. Cohen’s *d_z_*at the selected threshold was 1.40. **(C)** Mean paired ΔDSI, defined as late minus early DSI, plotted as a function of threshold *τ*. The dashed black line indicates the selected threshold (*τ* = 0.3), and the dotted red line indicates the threshold yielding the maximum mean paired ΔDSI (*τ* = 0.28). Mean paired ΔDSI at the selected threshold was 0.163. **(D)** Paired DSI values at the selected threshold (*τ* = 0.3) for all 20 nuclei pairs, with each pair corresponding to the same nucleus tracked from early to late chromatin reorganization. Blue points indicate early chromatin reorganization and orange points indicate late chromatin reorganization; gray lines connect paired measurements from the same nucleus. Mean ± SEM is overlaid for each group. DSI increased from the early to the late state of chromatin reorganization, and the paired difference was significant by Wilcoxon signed-rank test (*p* = 2.86 × 10^−6^).

**Supplementary Figure 2.**
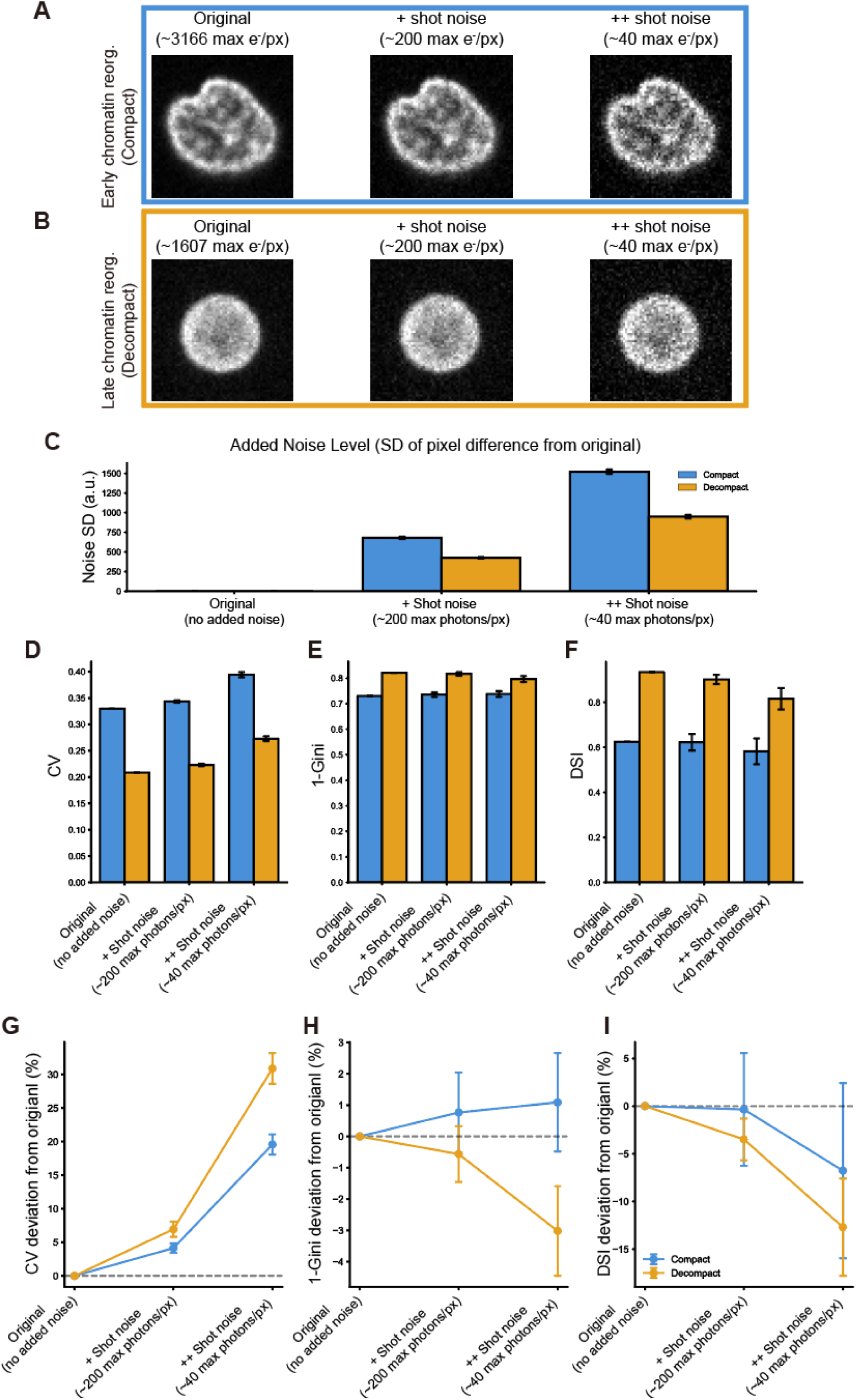
Effect of photon shot noise on CV, 1-Gini, and DSI. **(A, B)** Representative early-reorganization compact (A; original max ∼3,166 e−/pixel) and late-reorganization decompact (B; original max ∼1,607 e−/pixel) nuclei at the original signal level and after Poisson shot-noise degradation to ∼200 and ∼40 maximum photons per pixel. The degraded conditions are defined by the simulated photon budget assigned to the brightest pixel, not by measured electrons. The original signal is expressed as estimated detected photoelectrons (e−), calculated from raw intensities using the camera conversion gain (0.23 e−/ADU) after background subtraction. Because quantum efficiency correction was not applied, we report detected photoelectrons rather than direct photon counts; converting these values to photons reaching the camera sensor would only increase the estimated photon number. **(C)** Added noise level, quantified as the standard deviation of the per-pixel difference between each shot-noise-degraded image and its original, for the compact (blue) and decompact (orange) states (mean ± SD across N = 500 simulations). **(D–F)** CV **(D)**, 1-Gini **(E)**, and DSI **(F)** for the compact (blue) and decompact (orange) states across noise conditions (mean ± SD across N = 500 simulations). The two states remain separated at every photon budget. **(G–I)** Signed deviation of CV **(G)**, 1-Gini **(H)**, and DSI **(I)** from the noise-free value as a function of photon budget, for the compact (blue) and decompact (orange) states. All three metrics are affected by shot noise, with larger deviations for the lower-signal decompact state—the intensity-dependent behavior expected from shot-noise statistics. At the ∼40-photon budget, the decompact-state deviation reaches approximately +31% (CV), −13% (DSI), and −3% (1-Gini); 1-Gini is the least noise-sensitive, CV the most, and DSI intermediate. None of the three metrics is invariant to shot noise, in contrast to CV-ICS. In this representative fixed-mask simulation, the noise-induced bias did not inflate the compact-versus-decompact difference, and its sign was preserved at every photon budget tested; the effect was metric-dependent, reducing the apparent separation for 1-Gini and DSI, whereas for CV both states shifted upward and the separation was largely preserved. For reference, the representative spinning disk images correspond to a mean of ∼980–1,340 detected electrons per stored pixel (∼240–340 after 2×2 binning), well above the ∼40-photon regime in which shot-noise distortion becomes significant.

